# Protein language models enable prediction of polyreactivity of monospecific, bispecific, and heavy-chain-only antibodies

**DOI:** 10.1101/2023.11.06.565888

**Authors:** Xin Yu, Kostika Vangjeli, Anusha Prakash, Meha Chhaya, Samantha J Stanley, Noah Cohen, Lili Huang

**Affiliations:** Biotherapeutics and Genetic Medicine, AbbVie Bioresearch Center, 100 Research Drive, Worcester, MA 01605

**Keywords:** deep learning, protein language model, transfer learning, antibody discovery, bispecific antibody, single-domain antibody, non-specificity, polyreactivity

## Abstract

Early assessment of antibody off-target binding is essential for mitigating developability risks such as fast clearance, reduced efficacy, toxicity, and immunogenicity. The baculovirus particle (BVP) binding assay has been widely utilized to evaluate polyreactivity of antibodies. As a complementary approach, computational prediction of polyreactivity is desirable for counter-screening antibodies from *in silico* discovery campaigns. However, there is a lack of such models. Herein, we present the development of an ensemble of three deep learning models based on two pan-protein foundational protein language models (ESM2 and ProtT5) and an antibody-specific protein language model (Antiberty). These models were trained in a transfer learning network to predict the outcomes in the BVP assay and the bovine serum albumin (BSA) binding assay which was developed as a complement to the BVP assay. The training was conducted on a large dataset of antibody sequences augmented with experimental conditions, which were collected through a highly efficient application system. The resulting models demonstrated robust performance on normal mAbs (monospecific with heavy and light chain), bispecific Abs, and single-domain Fc (VHH-Fc). Protein language models outperformed a model built using molecular descriptors calculated from AlphaFold 2 predicted structures. Embeddings from the antibody-specific and foundational protein language models resulted in similar performance. To our knowledge, this represents the first application of protein language models to predict assay data on bispecifics and VHH-Fcs. Our study yields valuable insights on building infrastructures to support machine learning activities and training models for critical assays in antibody discovery.

## Introduction

For monoclonal antibodies (mAbs), it has been recognized that there are two types of off-target binding activities: polyspecificity and polyreactivity^1^. Polyspecificity is thought to be driven by factors such as molecular mimicry of the antigens, plasticity of the complementarity determining region (CDR), and the presence of multiple, possibly overlapping, paratopes. On the other hand, polyreactivity is associated with the presence of excess positive charge or hydrophobicity in the variable region, leading to low-affinity binding with matrix or membrane proteins through general nonspecific chemical interactions. While polyspecific interactions can sometimes be identified and eliminated, polyreactivity is often of unknown origin, presenting a greater challenge in antibody development^2,3^. Failure to address polyreactivity could lead to impact on pharmacokinetics (PK), potency, bioavailability, and immunogencity^1,2,4^.

Several binding assays have been developed to assess polyreactivity. These assays utilize HEK293 cells, double-stranded DNA (dsDNA), heparin, insulin, - or baculovirus particles (BVP)^2,4–7^. In the BVP assay, primary mAbs were added at high concentrations to BVP-coated plates and detected using a secondary mAb. This assay has been shown to correlate with clearance in humans and non-human primates^2^. A similar experiment can be done using bovine serum albumin (BSA) instead of BVP. While it is valuable to include these assays in early screening funnel, weak nonspecific interactions necessitate usage of high antibody concentrations, posing a challenge for small-scale/high-throughput protein production where material is scarce. As such, employing ML models to predict polyreactivity and guide the prioritization of clones for scale-up production would be useful. In addition, such models are critical for *in silico* antibody discovery campaigns as models trained on binding data alone lack the ability to predict specificity and therefore a non-specificity model is often required as a counter-screen method^8,9^.

Several such models for predicting antibody polyreactivity have been described, utilizing molecular descriptors from sequence and structures^6,10^, embeddings from one-hot residue representations^8,9^ or protein language models (PLMs)^11^. To our knowledge, no polyreactivity model has been developed using the BVP and BSA assays, nor has embedding from a PLM been used to predict functional properties of unconventional, bispecific formats. Herein, using a highly optimized data capture system, we generated a large BVP and BSA assay dataset augmented with experimental parameters. Using this dataset, we developed an ensemble of three deep learning models based on two PLMs (ESM2 and ProtT5) and one antibody-specific PLM (Antiberty). Our models demonstrated robust performance on polyreactivity prediction for normal mAbs (monospecific with heavy and light chain), single-domain Fc (VHH-Fc), and 12 subtypes of bispecific Abs.

## Results

### Development of an efficient application system to capture, analyze, visualize assay data and support machine learning activities

Obtaining high-quality data is the first and a crucial step in machine learning (ML). We designed and optimized an application system with four components: assay request, experiment planning, automated data analysis, and ML. The assay request component interfaces with Abbvie’s internal database for biologics registration, allowing tracking of metadata such as sequences and lot-specific concentrations. The experimental planning component captures user-defined experimental conditions and generates calculations. The data analysis component ingests and analyzes raw data from plate readers. While raw data (e.g. absorbance values from wells) is recorded in the database, secondary normalized data (e.g. fold over control) is computed on demand by applying a user-customizable function across the entire database. This ensures that the readout is always computed in a consistent manner specific to the user’s requirement, minimizing noise for modeling. With a few clicks, the application automatically generates interactive figures, Graphpad Prism files and PowerPoint reports, reducing a typical 3-hour analysis to under 15 minutes. In the ML component, we optimized model architecture, trained the models on existing data, evaluated them on new data and then fine-tuned them for the next cycle. A diagram of the application architecture, database schema, and snapshots of the application are shown in Figure 1. Unlike commercial data capture software, which often take many years to integrate with enterprise systems and optimize features, our application was built from scratch and deployed in about five months, representing a 10 to 30-fold faster development timeline compared to similar tasks. This underscores the importance of efficient application development to support ML activities.

**Figure 1.**
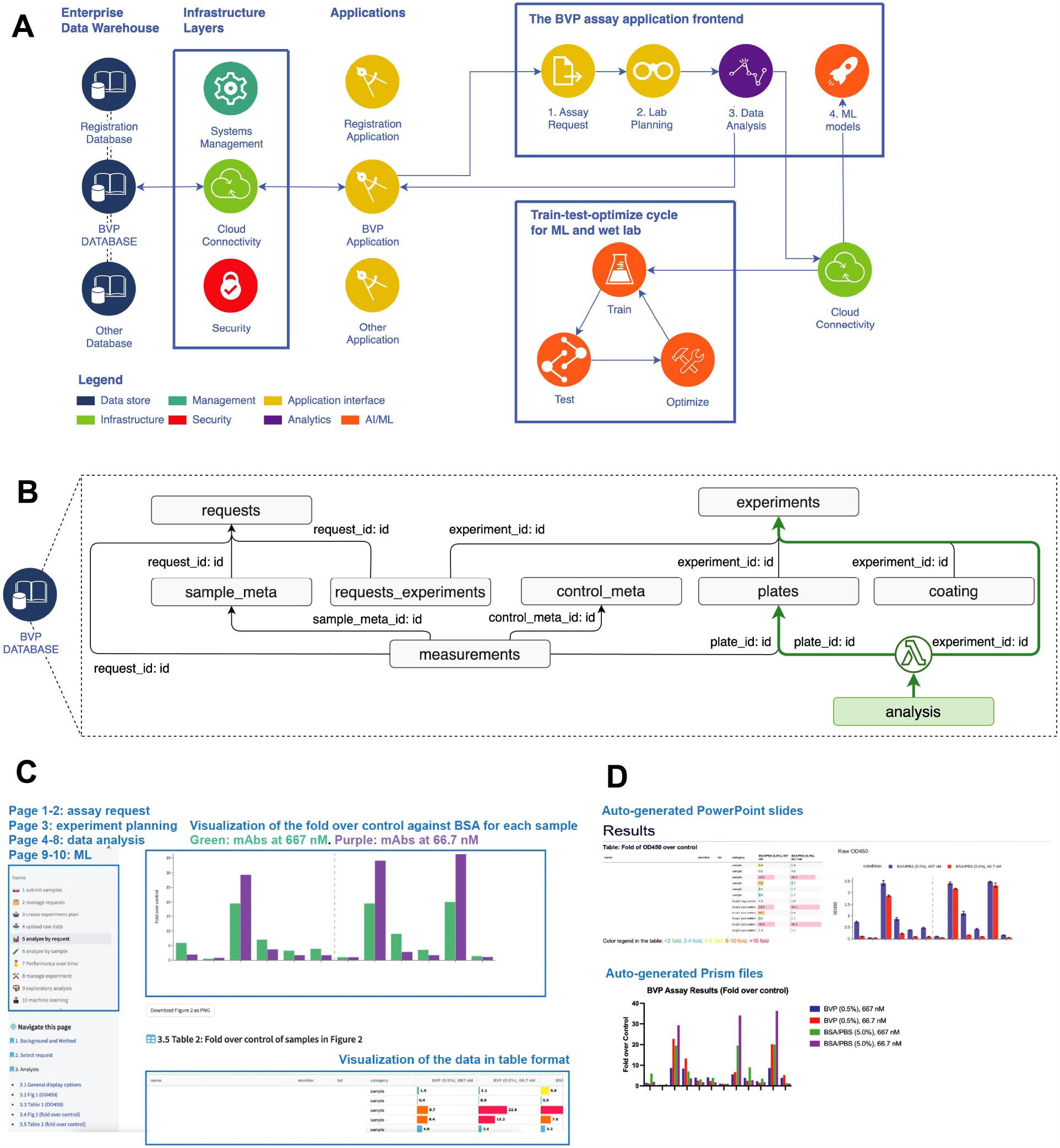
Overview of the application built to support data capture, analysis, visualization, and ML activities. **A**: Overview of the application system. Color of the icons indicates different types of data systems. **B**: Database schema. Primary and foreign key pairs were indicated on the arrows between tables. A Lambda function (green round icon) was deployed to enforce a real-time and consistent method of calculating the fold over control across the entire database, including legacy data. **C**: Snapshot of the application. The left side bar lists the pages. The bar plot is automatically generated by the application when raw data is uploaded. The lower table presents an alternative way of visualizing the data. **D**: The application automatically creates powerpoint slides and prism files.

### Exploratory data analysis

#### Description of the July dataset

The July dataset consisted of human IgG mAbs mostly in the discovery stage from our internal campaigns. We defined uniqueness on two levels. The unique mAbs refer to distinct sequences. Among the 509 unique mAbs, 81% had both heavy chain (HC) and light chain (LC), while the rest were HC only. All HC only mAbs were VHH-Fc. In addition, 86% were monospecific, with the remaining being bispecific. Of the bispecific mAbs, 50% were in the scFv Ig format, while the others adopted various formats (Figure 2A through 2D). The median sequence similarity scores were 0.5 for region var1, 0.7 for region var2, 0.3 for region var3, and 0.6 for region var4, suggesting that the sequences were very diverse, as shown in Figure 2E.

**Figure 2.**
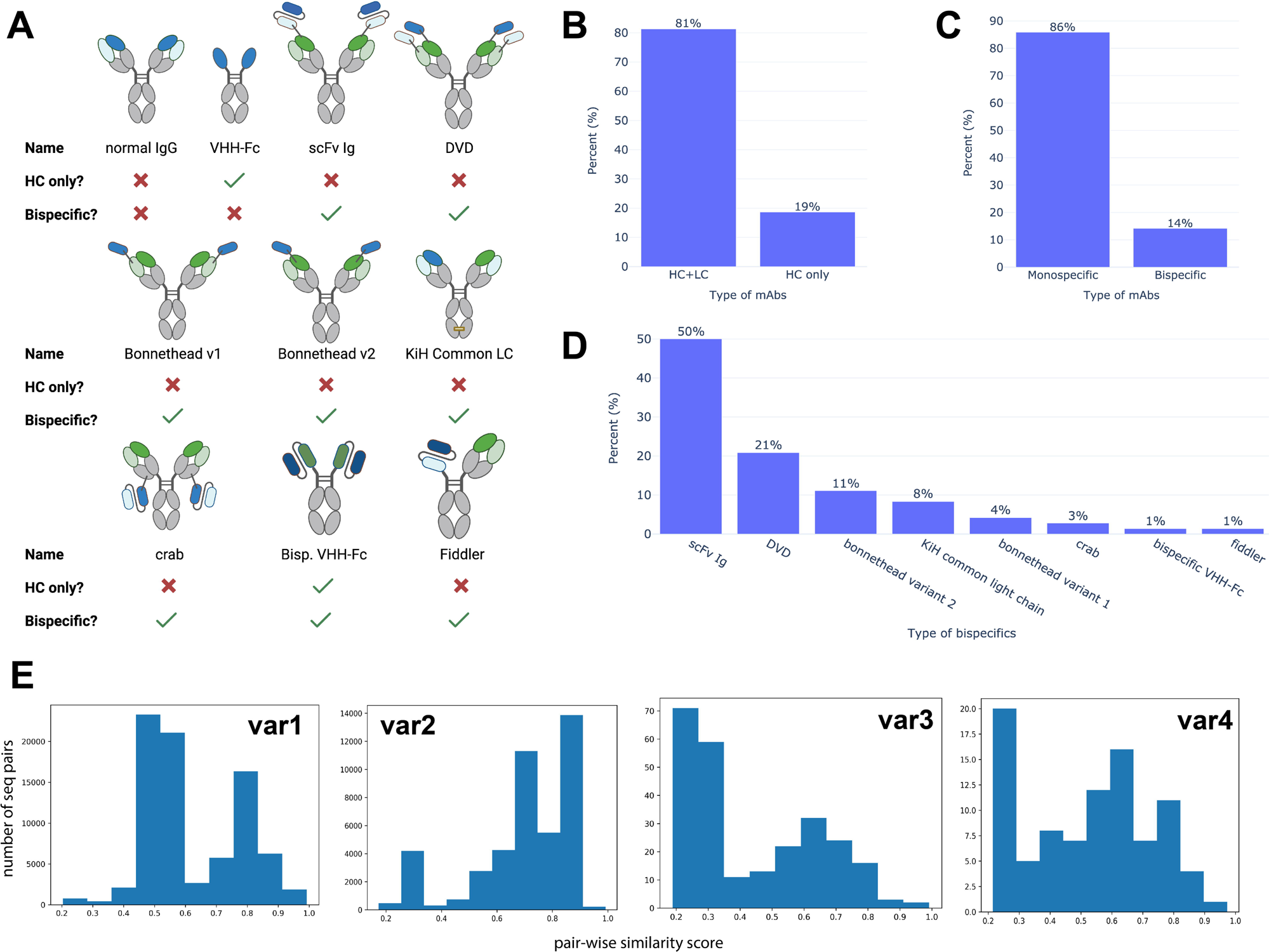
Distribution of N = 509 unique mAbs in the July dataset. **A**: Schematic drawing of mAb types. **B**: Percentage of HC-only (VHH-Fc) mAbs or mAbs with both HC and LC. **C**: Percentage of the monospecific (including monospecific normal mAbs and monospecific VHH-Fc) or bispecific mAbs **D**: Distribution of different bispecific formats, calculated as percentages of the total number of bispecific mAbs. E. Sequence similarity scores of variable regions var1, var2, var3, and var4. Definition of these regions were described in the method section where var i was calculated.

On the other hand, the unique data points refer to a unique combination of mAb sequence, concentration (ranging from 6.67 nM to 667 nM), and well coating type (either 0.5% BVP or 5% BSA). Unlike most literature models that omit experimental conditions, this approach had three benefits. First, it augmented the dataset from 509 unique mAbs to 1664 unique data points. Second, it reduced noise due to varying experimental conditions. Third, it gave the model an opportunity to learn about concentration-dependent effects of polyreactivity.

#### Concentration-dependent effects of polyreactivity

Polyreactivity assays often require high Ab concentrations, closer to *in vivo* concentrations when administered to humans. We assessed the impact of concentration on BVP and BSA ELISA by comparing the results of 313 unique mAbs tested at both 667 nM (roughly 100 µg/mL for a typical human IgG1) and 66.7 nM. Fold of signal over a negative control Ab were calculated. Based on data from a panel of positive control Abs with varying degree of polyreactivity and known to have clearance and/or toxicity issues, a threshold of 2 was set to distinguish between clean binders and polyreactive binders. Some antibodies showed polyreactivity at both 667 nM and 66.7 nM concentrations (Supp. Figure 1A), while some antibodies only appeared polyreactive at the higher concentration (Supp. Figure 1B). While not surprising, this result indicates that concentration needs to be taken into consideration when building models, if/when the dataset contains measurements at different concentrations.

#### Correlation of BVP and BSA ELISA

A positive correlation was observed between BVP and BSA ELISA for mAbs tested at 667 nM, with a Pearson correlation coefficient (R = 0.51), as shown in Figure 3.

**Figure 3.**
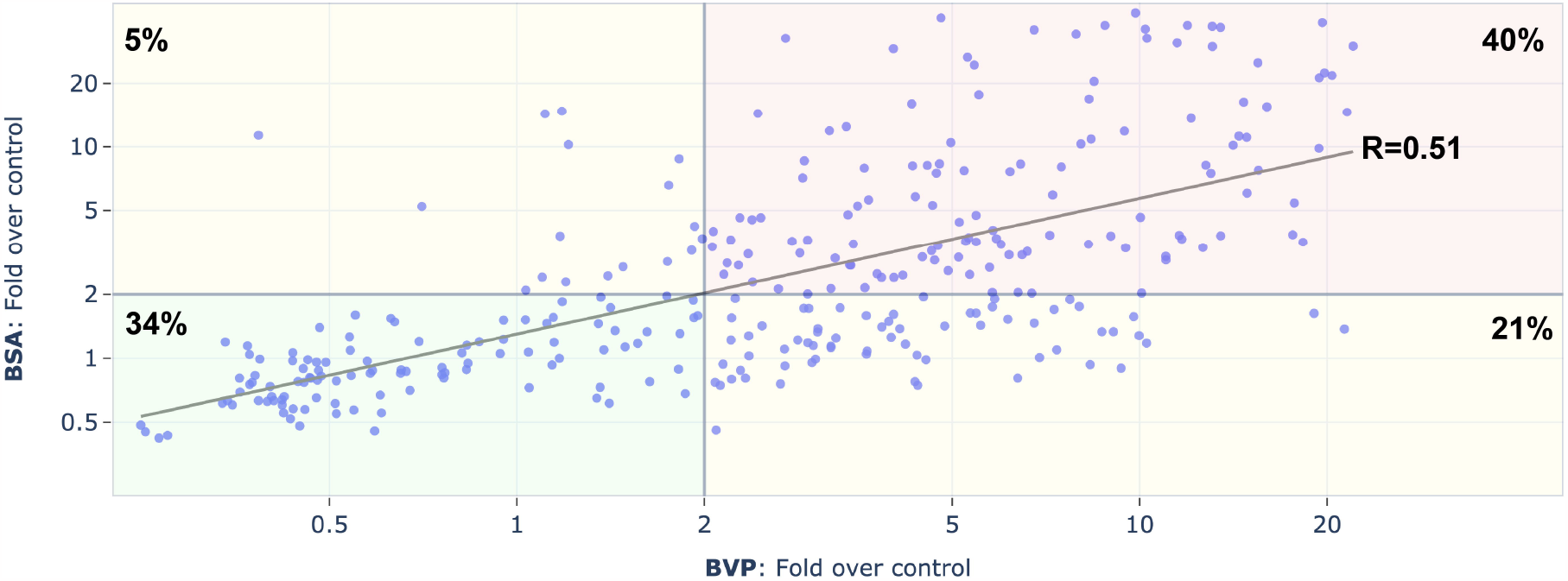
Correlation of BVP and BSA ELISA for 313 unique mAbs at 667 nM. Each dot is a unique mAb. The four quadrants categorize mAbs as clean (fold over control ≤ 2) or polyreactive (fold over control > 2) in one or both assays. Percentages of mAbs in each quadrant and the overall Pearson correlation coefficient were indicated on the figure.

Approximately 74% of mAbs were classified as either clean or polyreactive in both assays (lower left and upper right quadrants in Figure 3). Approximately 21% of mAbs showed polyreactivity on BVP but not on BSA, while 5% exhibited polyreactivity on BSA but not on BVP. The data indicates that polyreactivity detected by BVP and BSA ELISA have overlapping but different mechanisms.

#### Differences between allele families

We examined the relationship between polyreactivity and allele families of the HC and LC variable regions (a.k.a VH and VL, respectively). One-way ANOVA analysis on 228 unique monospecific mAbs (normal mAbs and VHH-Fc) tested at 667 nM in BVP and BSA ELISA showed statistical significance among VH families (p < 0.005).

Similar analysis on 134 monospecific mAbs (normal mAbs only) revealed significance among VL families (p < 0.005) (Supp. Figure 2). While some allele families (e.g.

IGHV1-69, IGKV1-27, IGKV1-9) seemed more prone to polyreactivity, the analysis was challenging to interpret. Possible reasons include the lack of consideration for heavy and light chain pairing, and some allele families having small sample sizes, both of which may confound the analysis. As a result, allele family information alone may not be very useful in predicting polyreactivity.

### Comparison of descriptor model and deep learning model

We compared two approaches for polyreactivity prediction: descriptor-based machine learning (a.k.a. the descriptor model) and PLM-based deep learning (a.k.a the PLM models). Both approaches utilized the July dataset augmented with experimental conditions. The details were described in the Method section, and a schematic drawing shown in Figure 4. Briefly, the descriptor model was built using AlphaFold multimer for VH/VL dimer structure prediction, followed by Schrodinger for sequence and structure descriptor calculation (Figure 4A). The PLM model 1 was trained using embeddings from Antiberty on a transfer learning network composed of 2-dimenional (2D) convolutional networks and classifier head (Figure 4, B through D).

**Figure 4.**
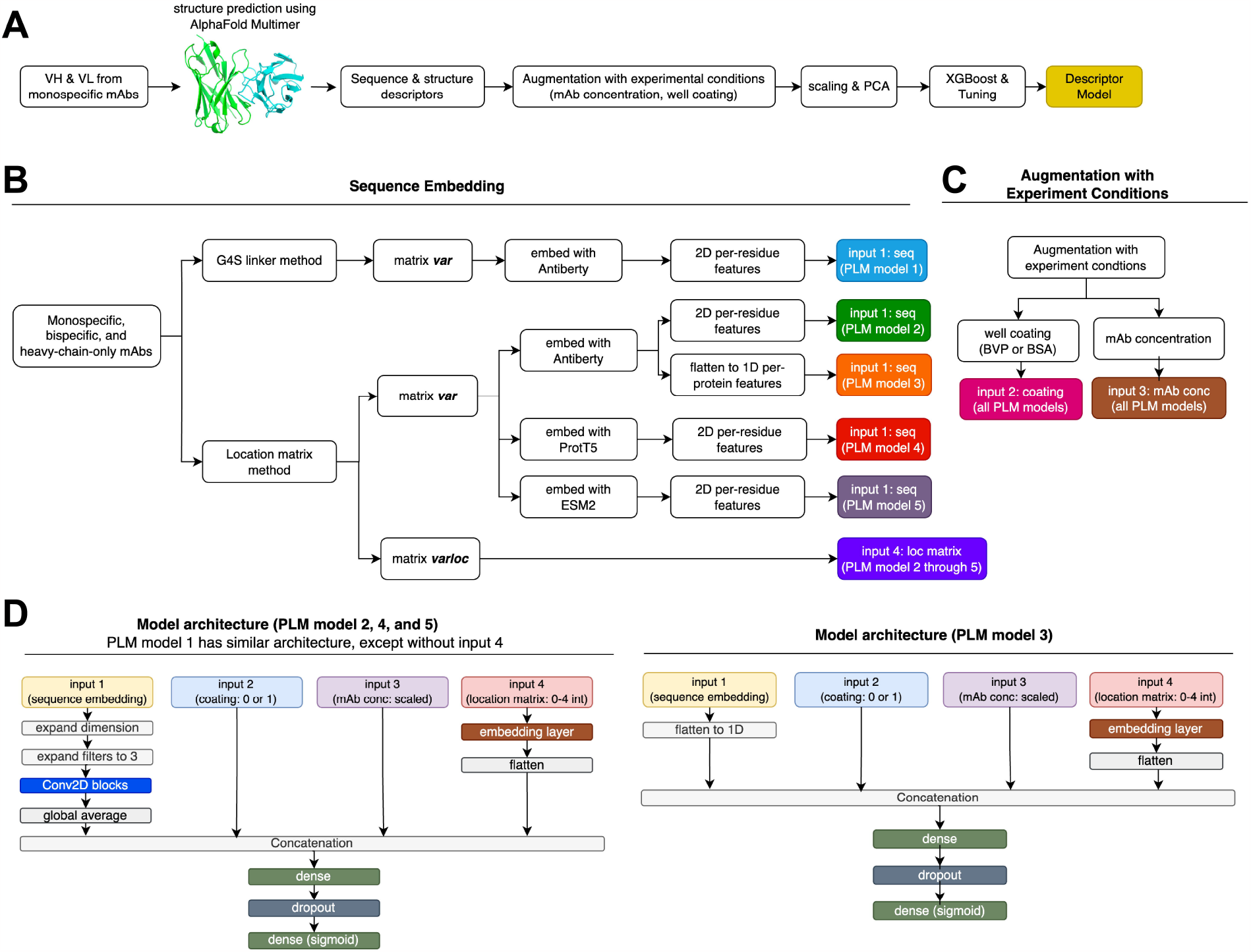
Design of the descriptor model and PLM models. **A**: VH/VL pairs from monospecific mAbs were modeled with AlphaFold, sequence and structure features (a.k.a. descriptors) computed using Schrodinger, and merged with experimental condition features. After scaling and principal component analysis (PCA), the data was used to train an XGBoost classifier. **B-D**: Components of the PLM models. **B**: Blocks labeled “input 1” in distinct colors represent different sequence embeddings generated by Antiberty (antibody-specific PLM), ProtT5 (pan-protein PLM), or ESM2 (pan-protein PLM), as either per-residue or per-protein features. Variable domain locations were encoded via either the G4S linker or the location matrix method. **C**: “input 2” and “input 3” represent well coating and mAb concentration, respectively. **D**: Transfer learning architecture of PLM model 1, 2, 4, and 5 (left) and 3 (right). The assembled network predicts polyreactivity as either clean (fold over control ≤ 2) or polyreactive (fold over control > 2).

Compared to the PLM model 1, descriptor model has several drawbacks. First, its application scope is limited to monospecific mAbs due to single VH/VL input constraint. In contrast, PLMs can handle bispecific mAbs by analyzing multiple variable region domains simultaneously. Second, the descriptor model has significantly lower throughput compared to PLMs. Prediction of a single VH/VL pair structure on AlphaFold takes 1-3 hours, while pretrained embeddings can be generated within seconds. Finally, when evaluated on blinded data, PLM model 1 (ROC AUC: 0.770 – 0.884) outperformed the descriptor model (ROC AUC: 0.711) in terms of area under the receiver operating characteristic curve (ROC AUC), as summarized in Table 1.

**Table 1.**
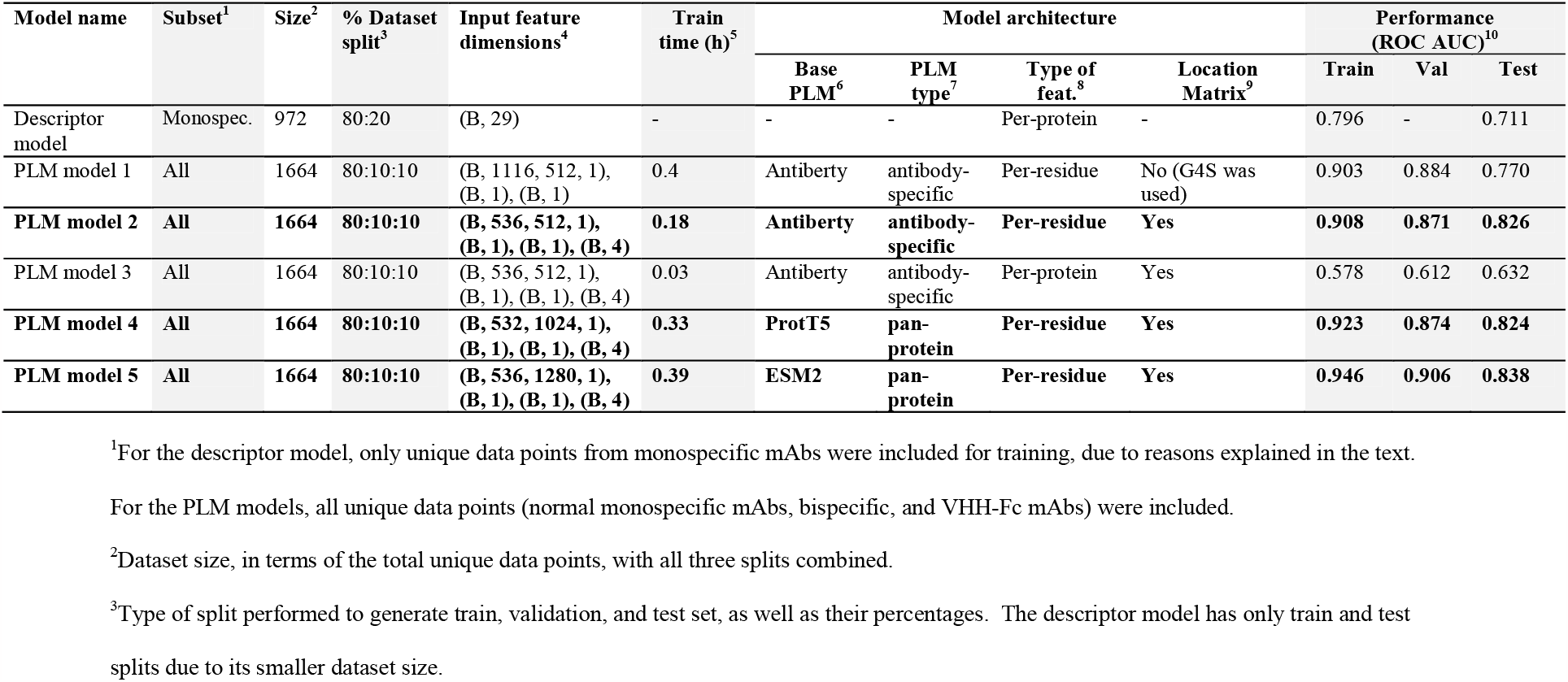
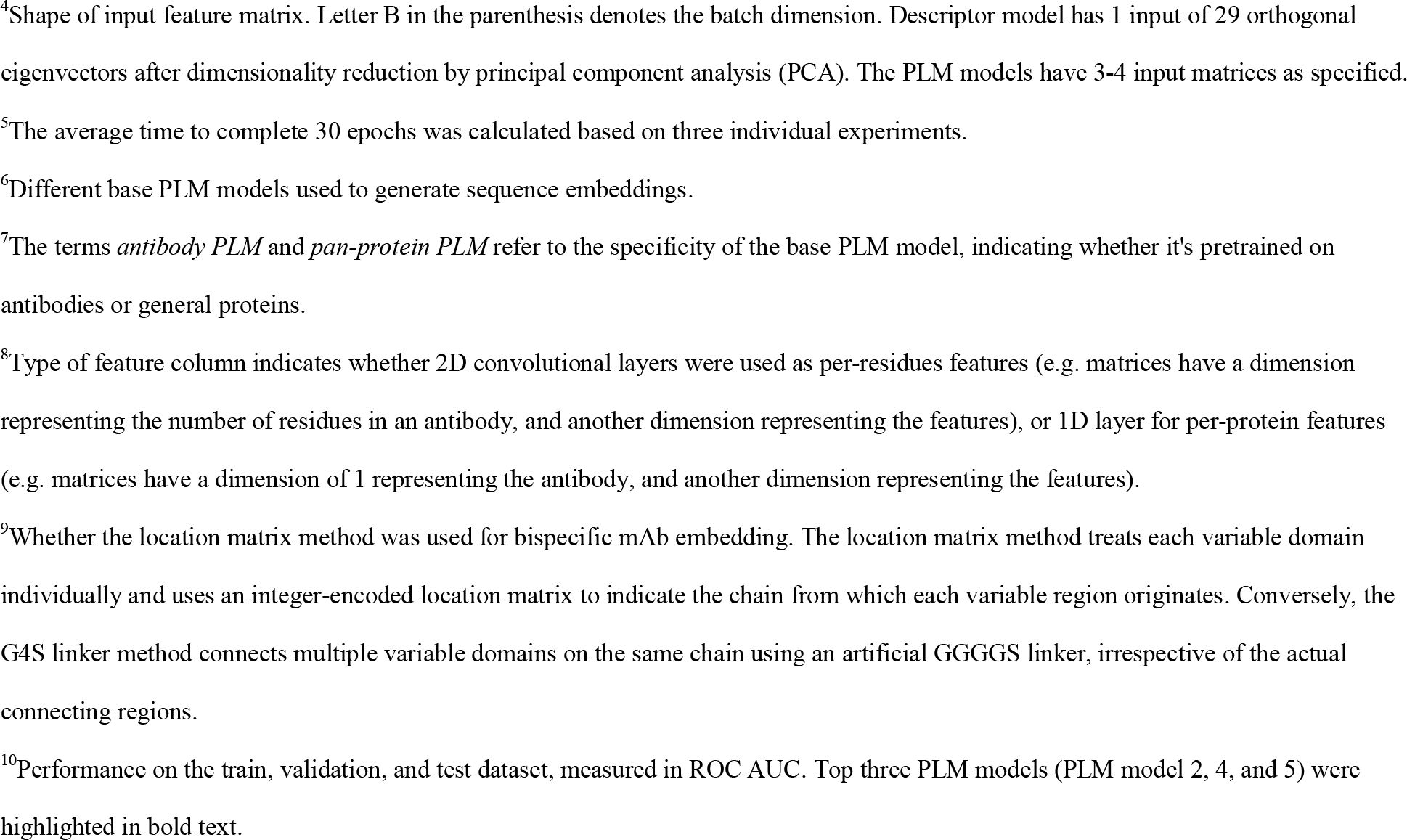
Summary of model performance in July dataset.

| Model name | Subset<sup>1</sup> | Size<sup>2</sup> | % Dataset split<sup>3</sup> | Input feature dimensions<sup>4</sup> | Train time (h)<sup>5</sup> | Model architecture |  |  |  | Performance (ROC AUC)<sup>10</sup> |  |  |
| --- | --- | --- | --- | --- | --- | --- | --- | --- | --- | --- | --- | --- |
|  |  |  |  |  |  | Base PLM<sup>6</sup> | PLM type<sup>7</sup> | Type of feat.<sup>8</sup> | Location Matrix<sup>9</sup> | Train | Val | Test |
| Descriptor model | Monospec. | 972 | 80:20 | (B, 29) | - | - | - | Per-protein | - | 0.796 | - | 0.711 |
| PLM model 1 | All | 1664 | 80:10:10 | (B, 1116, 512, 1), (B, 1), (B, 1) | 0.4 | Antiberty | antibody-specific | Per-residue | No (G4S was used) | 0.903 | 0.884 | 0.770 |
| **PLM model 2** | **All** | **1664** | **80:10:10** | **(B, 536, 512, 1), (B, 1), (B, 1), (B, 4)** | **0.18** | **Antiberty** | **antibody-specific** | **Per-residue** | **Yes** | **0.908** | **0.871** | **0.826** |
| PLM model 3 | All | 1664 | 80:10:10 | (B, 536, 512, 1), (B, 1), (B, 1), (B, 4) | 0.03 | Antiberty | antibody-specific | Per-protein | Yes | 0.578 | 0.612 | 0.632 |
| **PLM model 4** | **All** | **1664** | **80:10:10** | **(B, 532, 1024, 1), (B, 1), (B, 1), (B, 4)** | **0.33** | **ProtT5** | **pan-protein** | **Per-residue** | **Yes** | **0.923** | **0.874** | **0.824** |
| **PLM model 5** | **All** | **1664** | **80:10:10** | **(B, 536, 1280, 1), (B, 1), (B, 1), (B, 4)** | **0.39** | **ESM2** | **pan-protein** | **Per-residue** | **Yes** | **0.946** | **0.906** | **0.838** |

<sup>1</sup>For the descriptor model, only unique data points from monospecific mAbs were included for training, due to reasons explained in the text.
For the PLM models, all unique data points (normal monospecific mAbs, bispecific, and VHH-Fc mAbs) were included.
<sup>2</sup>Dataset size, in terms of the total unique data points, with all three splits combined.
<sup>3</sup>Type of split performed to generate train, validation, and test set, as well as their percentages. The descriptor model has only train and test splits due to its smaller dataset size.
<sup>4</sup>Shape of input feature matrix. Letter B in the parenthesis denotes the batch dimension. Descriptor model has 1 input of 29 orthogonal eigenvectors after dimensionality reduction by principal component analysis (PCA). The PLM models have 3-4 input matrices as specified.
<sup>5</sup>The average time to complete 30 epochs was calculated based on three individual experiments.
<sup>6</sup>Different base PLM models used to generate sequence embeddings.
<sup>7</sup>The terms *antibody PLM* and *pan-protein PLM* refer to the specificity of the base PLM model, indicating whether it's pretrained on antibodies or general proteins.
<sup>8</sup>Type of feature column indicates whether 2D convolutional layers were used as per-residues features (e.g. matrices have a dimension representing the number of residues in an antibody, and another dimension representing the features), or 1D layer for per-protein features (e.g. matrices have a dimension of 1 representing the antibody, and another dimension representing the features).
<sup>9</sup>Whether the location matrix method was used for bispecific mAb embedding. The location matrix method treats each variable domain individually and uses an integer-encoded location matrix to indicate the chain from which each variable region originates. Conversely, the G4S linker method connects multiple variable domains on the same chain using an artificial GGGGS linker, irrespective of the actual connecting regions.

### Optimization of PLM models

#### Embedding bispecific mAbs using G4S linker or location matrix

Embedding bispecific mAbs is challenging because of different arrangements of the variable domains. As shown in Figure 2A, in formats like scFv Ig, DVD, bonnetheads, bispecific VHH-Fc, and the scFv arm of Fiddler, variable domains are connected by short peptides. Conversely, in the crab format, the VL and single-chain variable fragment (scFv) are separated by the light chain constant region. Additionally, some formats have bispecific domains only on the HC (e.g., scFv Ig, bonnethead v2, bispecific VHH-Fc), some only on the LC (e.g., bonnethead v1, crab), and some on both chains (e.g., DVD, KiH common light chain, Fiddler). A brute-force approach to encode entire HC or LC is impractical, not only due to Antiberty’s 510-amino acid input limitation but also because the large embedding matrices from long sequences would require substantial computational resources for training.

To address this challenge, for PLM model 1 we extracted variable domains on the same HC or LC then concatenated them using a G4S linker (Figure 4B). This keeps the input matrix size manageable and retains some positional information. Specifically, the resulting input shapes are (B, 1116, 512, 1) for PLM embedding, and (B, 1) each for concentration and coating embeddings, where B is the batch dimension. While this setup uses the G4S linker to indicate whether two variable domains share the same chain, it lacks detail about which specific chain (e.g. HC1 or HC2) they originate from. In addition, models like Antiberty, ProtT5, and ESM2 are primarily trained on natural protein sequences. Since endogenous mAbs typically feature one variable domain per chain, embedding multiple domains in a single sequence potentially raises questions about suitability.

PLM model 2 uses an integer-encoded location matrix to identify each variable domain’s origin chain (Figure 4B). By performing separate embedding on each variable domain, it aligns more closely with training data of Antiberty. In addition, reducing the input feature dimension cut the training time from 0.4 to 0.18 hours. Although it slightly lowered validation accuracy from 0.884 to 0.871 (1% decrease), it boosted test AUC from 0.770 to 0.826 (7% increase), prompting us to continue with this approach (Table 1).

#### 2D Per-residue feature vs 1D per-protein feature

Both PLM models 1 and 2 use 2-dimensional (2D) convolutional neural networks, treating per-residue features as images similar to the approach described in our previous study^12^. An alternative approach, PLM model 3, flattened these into 1D per-protein features, similar to the descriptor model but without explicit association with sequence and structure features (Figure 4B). PLM model 3 underperformed across all dataset splits (Table 1), showing significantly worse performance (up to 36% decrease in AUC) compared to PLM models 1 and 2 as well as the descriptor model. This highlights the importance of per-residue features and convolutional networks in our case.

#### Antibody-specific PLM vs pan-protein foundational PLMs

Switching from antibody-specific PLM Antiberty to pan-protein foundational PLMs like ProtT5 and ESM2 resulted in similar performance. PLM model 4, which utilized ProtT5, had an AUC of 0.874 on the validation set and 0.824 on the test set. PLM model 5, which was built from ESM2, yield an AUC of 0.906 on the validation set and 0.838 on the test set. This suggests that when used as embeddings, PLMs pretrained on antibody sequences might not necessarily offer a significant advantage over those pretrained on general proteins.

### Performance of PLM model 5 on subsets of monospecific, bispecific, and VHH-Fc

Using the July datasets, we further assessed performance of PLM model 5 on the combined validation and test sets which the model was not trained on. The analysis in Figure 5 (A through C) provided a breakdown of its efficacy across different types of mAbs. The overall AUC was 0.871, with an AUC of 0.884 for mAbs containing both heavy and light chains, compared to 0.802 for VHH-Fc. The model performed robustly on bispecific mAbs (AUC 0.906) and monospecific mAbs (AUC 0.865).

**Figure 5.**
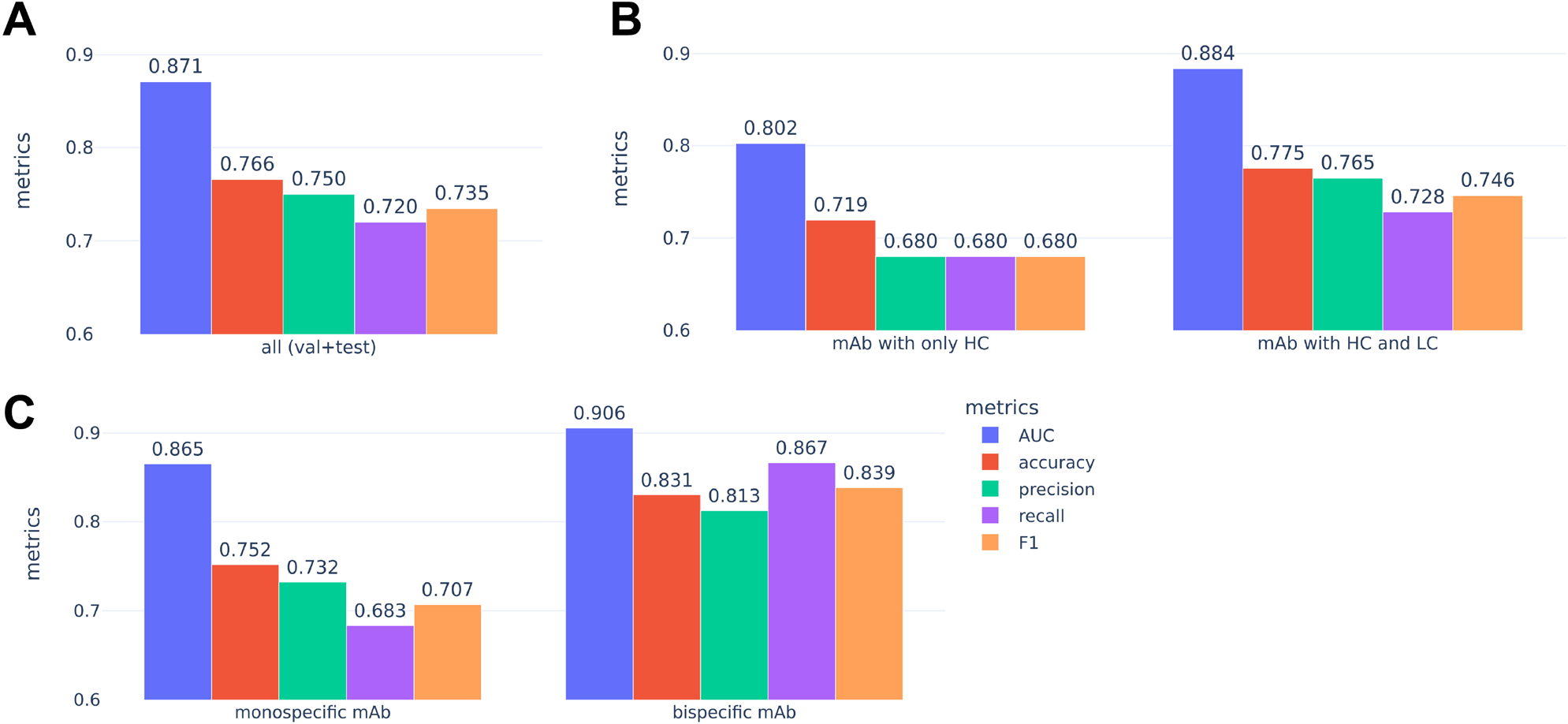
Performance of PLM model 5 on the combined validation and test sets (N=333 blinded data points) from the July dataset. A panel of metrics were shown in different colors. **A**: Overall performance. **B:** performance on HC-only mAbs (VHH-Fc, N = 57) and mAbs with both HC and LC (N=276). **C**: Performance on monospecific (N=274) and bispecific mAbs (N=59). Monospecific mAbs include both normal IgG as well as monospecific VHH-Fc.

### Confirmation of findings with the September dataset

#### Description of the September dataset

Our application collected 321 unique mAbs during a period of approximately 2 months, including 282 that were not presented in the July dataset. This new batch differed notably from the original July dataset. For example, 99% had both HC and LC (Figure 6A). About 23% were bispecific, including four new undisclosed subtypes (Figure 6C). Due to these differences, the new data was combined with the original July dataset to form the September dataset, totaling 2,998 unique data points augmented with experimental conditions. The median sequence similarity score for var1, var2, var3, and var4 as well as distribution of these scores were similar to the July dataset.

**Figure 6:**
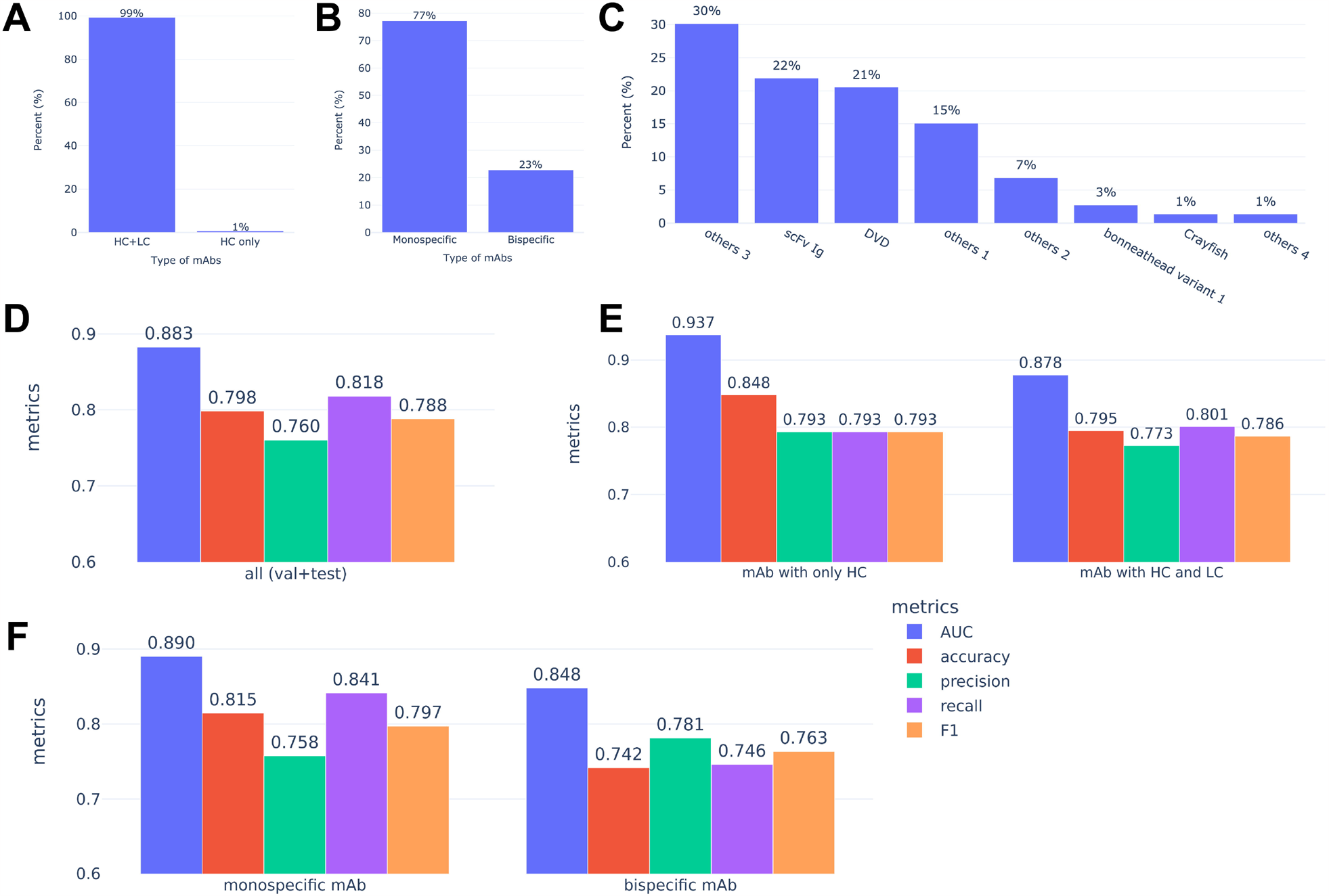
Description of the new data in the September dataset and the performance of the ensemble model. **A-C**: Distribution of N = 321 unique new mAbs added during a 2-month period. **A**: Percentage of HC-only (VHH-Fc) mAbs or mAbs with both HC and LC. **B**: Percentage of the monospecific or bispecific mAbs. Monospecific mAbs include both normal IgG and monospecific VHH-Fc. **C**: Distribution of different bispecific formats. Others 1 through 4 refer to undisclosed bispecific formats that did not exist in the July dataset. **D-F**: Performance of the ensemble model built from PLM 2, 4, and 5 on the combined validation and test sets (N=600 blinded data points) from the September dataset. **D**: Overall performance. **E**: Performance on the HC-only mAbs (N=79) and mAbs with both HC and LC (N=521). **F**: Performance on monospecific (N=480) and bispecific mAbs (N=120).

#### Performance of the PLM models and the ensemble

We fine-tuned PLM models 2 (Antiberty-based), 4 (ProtT5-based), and 5 (ESM2-based) on the September dataset using a similar approach as before. All models showed robust overall performance, with AUCs ranging from 0.862 to 0.891 on unseen data in the validation and test sets (Table 2). This again confirmed embeddings from antibody-specific PLMs performed comparably to those from foundational PLMs. Each model also maintained strong performance (AUC 0.812–0.968) across different mAb categories: monospecific, bispecific, and VHH-Fc (Supp. Table 1). Ensemble of PLM models 2, 4, and 5 further improved some of these metrics, with final AUC values ranging from 0.848 to 0.937 across different mAb formats (Figure 6D through F).

**Table 2.** Summary of model performance in September dataset^1^.

| Model name | Subset | Size | % Dataset split | Train time (h) | Base PLM | Performance in (ROC AUC) |  |  |
| --- | --- | --- | --- | --- | --- | --- | --- | --- |
|  |  |  |  |  |  | Train | Val | Test |
| PLM model 2 | All | 2998 | 80:10:10 | 0.3 | Antiberty | 0.950 | 0.883 | 0.862 |
| PLM model 4 | All | 2998 | 80:10:10 | 0.54 | ProtT5 | 0.933 | 0.891 | 0.862 |
| PLM model 5 | All | 2998 | 80:10:10 | 0.66 | ESM2 | 0.901 | 0.878 | 0.863 |

<sup>1</sup>Column interpretations are consistent with those in Table 1.

### Mechanistic insights

Prior studies have reported the important role of hydrophobicity^1,13,14^ and charge^1,6,14,15^ in polyreactivity. In BSA ELISA, the detected polyreactivity of mAbs is likely driven by charge, since BSA has an isoelectric point (pI) of around 5 and is therefore negatively charged in pH 7.4 buffer. Data from BSA assay correlates well with that from other negatively charged ELISA such as dsDNA assay (data not shown). The GP64 trimer envelope protein of BVP^16,17^, modeled via AlphaFold, displayed hydrophobic regions along its length. The tip of the trimer presented an area of strong hydrophobic surface (Supp. Figure 3). These regions could potentially be associated with hydrophobicity-driven polyreactivity. In addition, wells in BVP ELISA were coated with BVP but blocked with BSA. It is therefore conceivable that the detected polyreactivity in BVP ELISA could be likely driven by hydrophobicity and/or charge. These observations are corroborated by the SHapley Additive exPlanations (SHAP) scores from the descriptor model. According to the SHAP scores, the top contributing features were mAb concentration, hydrophobicity (LC CDR1), well coating, and positive charges (HC CDR1 and LC FR3, Supp. Figure 3). Notably, the highest SHAP score (0.519) was attributed to mAb concentration, confirming the importance of this factor in predictive modeling.

## Discussion

It is well established that high-quality data is crucial for machine learning activities^18^. High-quality data improves the model performance and facilitates the resolution of complex tasks. In contrast, noisy, irrelevant, or erroneous data can often mislead algorithms. Unfortunately, in pharmaceutical industry, a domain rich with historical and continually generated data, there is currently a lack of methods to efficiently integrate data capture with machine learning activities. Herein, we demonstrated an example where assay design, lab automation, data capture, analysis, visualization, and machine learning were orchestrated as a cohesive system. Measures such as enforcing duplicate wells of measurements, utilization of liquid handler, and normalization of signal using a standard control mAb were implemented to mitigate the impact of manual pipetting errors and fluctuations of raw absorbance values in the assay. In practice, historical data and user preferences sometimes demand flexible normalization against different control mAbs. This poses potential risks for data inconsistency in conventional data capture software because the fold over control value is often stored as a single field without metadata on how the experiment was conducted and analyzed. To address this, we captured experimental conditions like well coating and mAb concentration, enabling consistent real-time data aggregation and calculations of fold over control. This ensures data quality while allowing user and project-specific interpretations. Additionally, the inclusion of experimental metadata as input features for the models was proven to be critical for model performance, as we demonstrated by SHAP score. This aligns well with the current trend in multimodal machine learning^19^, where different modalities of data (e.g. sequence, structure, experimental metadata, text and image) would eventually be combined to develop useful models for real-world applications.

We explored whether an antibody-specific PLM is always preferable over a foundational PLM, for antibody related tasks. Literature has shown that pretraining PLMs on antibodies can enhance model performance. For instance, IgFold, which uses Antiberty, predicts antibody structures with similar or better accuracy, but faster than, AlphaFold^20^. On the other hand, foundational PLMs have also proven effective in affinity maturation of antibodies, indicating their generalizability across protein families^21^. Some researchers suggested that while the ability of antibody-specific PLMs to address CDR’s hypervariability is advantageous, their smaller, less diverse training set limits the depth of insights that could be otherwise obtained in foundational PLM^22^. In our study, Antiberty and foundational PLMs showed comparable performance in polyreactivity datasets. This could be possibly due to polyreactivity being driven by low-affinity, nonspecific chemistry due to hydrophobicity and charged patches in the variable region, rather than high-affinity, antigen-specific interactions within the CDRs. This highlights the potential value in comparing and possibly integrating both PLM types to leverage their respective strengths for future problems.

There are several merits of our study. First, it pioneers the application of PLM embeddings to diverse mAb formats like monospecific, bispecific (12 subtypes), and VHH-Fc. By demonstrating uniform embeddings and simultaneous training across these mAb types, we took a step to address the growing interest in developing such formats as therapeutics or tool reagents. Second, we directly compared descriptor model and PLM models, illustrating their complementary nature. The PLM models had a broader application scope and achieved better performance. On the other hand, the descriptor model yielded mechanistic insights that is consistent with empirical observations (e.g. concentration dependent effect) and literature data^1^ (e.g. hydrophobicity and charge as key factors driven polyreactivity). Third, when bioassay data was involved for modeling, previous studies in the literature often relied on a one-time collection of datasets from the same or different sources^5,11,23^. One-time data collection leaves open the question about whether the conclusions hold as the new data comes in. The different data sources raise the possibility that data from different assays and experimental conditions might be inconsistent. We tackled the issue by building an application that integrated key steps from data capture to machine learning. We were able to repeatedly tune the models on datasets that were collected overtime, under the same experimental conditions. This provides us an opportunity to test the previous conclusions and derive new insights.

There are several caveats of our models. First, our current model handles up to 4 variable regions in a single mAb. Second, in either the G4S linker method or the location matrix method, the connection region between the variable domains is lost.

Because these regions could be artificial linkers of different lengths, or constant regions of the heavy or light chain, they contain important spatial and steric information. Third, the Fc region was not included in the embedding due to its general similarity and the input length restriction from the PLMs. The impact of mutations in the human IgG1 Fc region on BVP/BSA ELISA remains unclear. Positive charges in the variable domain, which could drive polyreactivity detectable by BVP/BSA ELISA, has been shown to directly affect FcRn-dependent PK^1,6^. Interplay of these factors (charge, hydrophobicity, and Fc mutations) on the polyreactivity and PK requires further investigation. Fourth, currently the PLMs do not encode post-translational modification (PTM) features.

PTMs like sulfation and glycosation can alter antibody structure and paratopes, therefore potentially introducing changes to polyreactivity. Finally, the feature matrix is still far from complete. Factors absent from experiment metadata such as production cell line, protein purity, buffer composition, and freeze-thaw cycle could affect assay readout. These factors have not yet been captured as structured data. In additions, assays that are related to hydrophobicity (e..g retention time in hydrophobic interaction chromatography a.k.a HIC) and stability (e.g. affinity-catpure self-interaction nanoparticle spectroscopy a.k.a. AC-SINS, dynamic light scattering a.k.a. DLS) might be useful input features for polyreactivity models. If these dots are connected, we could then move closer towards multi-modal machine learning.

In conclusion, we developed an ensemble of models based on antibody-specific and foundational PLM embeddings to predict the polyreactivity in mAbs. These models can be potentially used as counter selection in combination with affinity models for in silico screening of antibodies with desired properties. Our study yields valuable insights on building infrastructures to support machine learning activities and training deep learning models for critical assays in antibody discovery.

### Materials and Methods

### Development data application

The database was built with PostgreSQL, and the application was developed using python web framework Streamlit^24^, then deployed on Amazon Web Services (AWS). Cloud connectivity and Lambda functions were established using either AWS or Cloudera.

### BVP and BSA ELISA

BVP ELISA was adapted from Hotzel, et al^2^. Negative charge based BSA ELISA was developed as a complementary assay to BVP assay and implemented conveniently in the same high throughput experiment. The assays were extensively optimized for automation on EP Motion liquid handler (Eppendorf). Briefly, BVP particles (LakePharma, Cat# SR-17000) were coated on half of a 384-well plate (Greiner Bio, Cat# 781061) at a 0.5% concentration in 50mM sodium bicarbonate buffer (Pierce, Cat# 28382). The other half of the plate were coated with just the sodium bicarbonate buffer. After overnight incubation at 4°C, plates were washed with 1x TPBS (1x PBS with 0.05% Tween 20) and blocked with 5% BSA in 1xPBS for 1 h at room temperature (RT). mAbs were added at 25 μl/well in duplicates to the BVP and BSA coated wells.

While the standard mAb concentrations were 667 nM and 66.7 nM, when materials were scarce, adjustments were made to these concentrations, resulting in a range from 6.67 nM to 667 nM in our datasets. Plates were incubated for an hour at RT, washed three times, and secondary antibody (Thermo, Cat# 31412) was added at a 1:10000 dilution in 1% BSA in 1x PBS. After 45 min of incubation at RT, plates were washed, and TMB substrate (Thermo, Cat# 34029) were added at 25 μl/well and incubated for 3 min at RT. Stop solution (VWR, Cat# BDH7500-1) were added at 25 μl/well. Readings were taken on a Clariostar Plus microplate reader (BMG Labtech). A low-polyreactive human IgG1 served as the normalization control mAb in each experiment. Given a specific mAb concentration and type of coating, fold over control was calculated as the ratio of the average signal of the sample mAb to that of the control mAb.

### Acquisition of the July dataset and exploratory data analysis

The July dataset was compiled using the previously described application, collecting data from internal BVP and BSA assays that were conducted primarily for mAbs in the discovery stage. Using Python packages abnumber (v0.3.2) and anarci (v2020.04.23), VH and VL domains were extracted from IgG HC and LC to calculate number of variable domains, chain stoichiometry, and germline families. This data was used to classify mAbs as monospecific, bispecific, or heavy-chain-only (VHH-Fc). Subtypes of bispecific mAbs (e.g. scFv-Ig, DVD, etc) were manually annotated. Statistical analysis was generated using scipy (v1.11.1), and figures generated using plotly (v5.16.1) and matplotlib (v3.7.1). Notably, a “unique mAb” is defined by its unique sequence and was utilized in analysis such as mAb type distribution, BVP and BSA ELISA correlation, and germline family data correlation. Conversely, “unique data point” is defined by a unique mAb sequence combined with mAb concentration and well coating. This was used for analyzing concentration effects and training descriptor and PLM models with experimentally augmented datasets.

### Training the descriptor model on the July dataset

A total of 972 unique data points on monospecific mAbs from the July dataset were utilized for building the descriptor model. These mAbs were modeled as VH/VL heterodimers using AlphaFold 2 (v2.3.0), with the top-ranked relaxed structures selected for 910-descriptor computation in Schrodinger Maestro (v2022-3). The resulting feature matrix was combined with experimental data (mAb concentration and coating), standardized, and underwent principal component analysis (PCA), yielding 29 orthogonal vectors for XGBoost input. Parameter tuning involved 500 trials on an 80/20 train-test split, and performance was assessed via 5-fold cross-validated (5CV) ROC AUC scores. A diagram of the process is shown in Figure 4A.

### Training the PLM models on the July dataset

A total of 1664 unique data points on monospecific, bispecific, and heavy-chain-only mAbs from the July dataset were utilized for building the PLM model 1 through 5. The sequences were processed taking stoichiometry into account. Specifically, let *HC*1,*LC*1,*HC*2,*LC*2 be the single amino acid sequence of heavy chain 1, light chain 1, heavy chain 2, light chain 2 sequences of a mAb. All HC sequences in our dataset satisfy the following constraint:

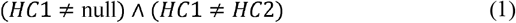

All LC sequences satisfy the following constraint:

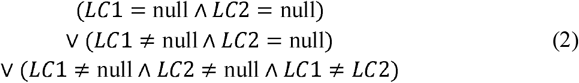

For example, a monospecific mAb and a bispecific DVD have non-null *HC*1 and *LC*1, but null *HC*2 and *LC*2. A monospecific VHH-Fc mAb have non-null Hc1 and null values for *LC*1,*HC*2 and *LC*2. Define *f*(*x*) as a function from abnumer or anarchi to extract all variable domains (VH and VL) from the input sequence *x*. Then:

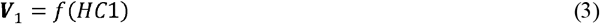

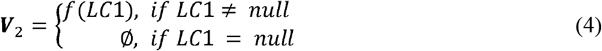

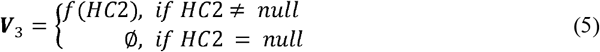

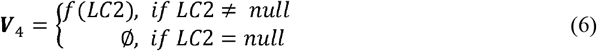

*V*_*i*_ where *i* ∈ {1, 2,3,4} is a set of VH and/or VL domains in chain *i*, arranged in the order they appear from the N-terminus to the C-terminus on that chain.

Let *var*_*i*_ where *i* ∈ {1, 2,3, 4} denote the arbitrarily defined 4 sequences used to construct the input matrix ***var*** for PLM embedding. For the G4S linker method, let *g*(***Y***) denote a function that concatenates the variable domains in set ***Y*** to a single sequence by connecting them with a short GGGGS peptide, in the order they appear. Then:

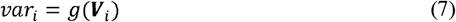

In the location matrix method, let ⨁ denote the ordered concatenation of multiple vectors into a single vector, without intermixing the elements within each vector. Let ***A*** denote the concatenated vector:

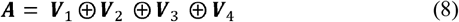

The current approach allows up to 4 variable domains per antibody. That is, we are guaranteed that with

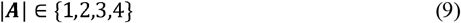

We can assign values to *var*_*i*_ as:

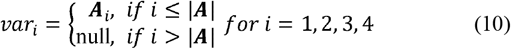

Let *varloc*_*i*_ denote the chain location of *var*_*i*_. Encode the location matrix *varloc* with:

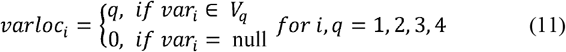

Sequences in ***var*** were then fed into PLM base models Antiberty^25,26^, Prot5_XL_Uniref^26^, ESM2^27^ to generate per-residue embeddings, as per default guidelines in their github repositories. The concentration matrix was normalized to a number between 0 and 1 using the following formula:

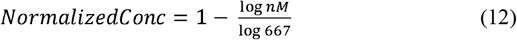

Where *nM*is the original concentration in nM. The coating matrix was encoded 0 if 5% BSA was coated on wells and 1 if 0.5% BVP was coated. The label matrix was encoded 0 if signal over control mAb was less than or equal to 2, and 1 otherwise. These matrices were fed into a deep learning model similar to our previous study^12^, as depicted in Supp. Figure 4. An 80/10/10 random split was used to generate train, validation, and test sets. The model was trained only on the train set, using an Adam optimizer with custom learning rate scheduler consisting of 10 epochs of linear warmup, 10 epochs of constant rate at 1e-3, and 10 epochs of cosine decay. The ROC AUC, accuracy precision, recall, and F1 metrics were calculated using sklearn package (v1.3.0) and plotted using Plotly (v5.14.1). A sketch of this process is shown in Figure 4 (B through E). Example of training script using Antiberty embedding is provided in as Supplementary Code.

### Building the ensemble model on the September dataset

The September dataset was formed by combining the July dataset with new entries collected over a 2-month period. We applied an 80/10/10 train-validation-test split, blinding the validation and test sets during tuning. Individual PLM models 2, 4, and 5 were first individually tuned, then combined into an ensemble using soft voting. The highest-AUC model received a 0.5 weight, and others 0.25. Details of training and evaluation process were consistent with prior methods.

### Mechanistic studies

Feature importance from the descriptor model was calculated using the SHapley Additive exPlanations (SHAP) scores^28^. GP64 trimer envelope protein of BVP was modeled using AlphaFold multimer.

## Abbreviations

1D: 1 Dimensional
2D: 2 Dimensional
Ab: Antibody
AC-SINS: Affinity-Capture Self-Interaction Nanoparticle Spectroscopy
ANOVA: Analysis of Variance
AUC: Area under the Curve
BSA: Bovine Serum Albumin
BVP: Baculovirus Particle
CDR: Complementarity Determining Region
DLS: Dynamic Light Scattering
ELISA: Enzyme-Linked Immunosorbent Assay
FcRn: neonatal Fc receptor
FR: Framework Region
HC: Heavy Chain
HIC: Hydrophobic Interaction Chromatography
LC: Light Chain
mAbs: Monoclonal Antibodies
ML: Machine Learning
PK: Pharmacokinetics
PLM: Protein Language Model
PTM: post-translational modification
ROC: Receiver Operating Characteristic curve
SHAP: SHapley Additive exPlanations
VH: Heavy Chain Variable Region
VHH-Fc: Single-Domain Fc
VL: Light Chain Variable Region

## Acknowledgement

The authors would like to thank the following AbbVie colleagues: Emma Fung, Gautam Sahu, Supriya Jalimarada-Shivakumar, Renee Miller, Olha Nazarko, Yifeng Lu, and Kangwen Deng for their valuable inputs and constructive feedback on the assay design, optimization, and writing; Andrew Goodearl for his scientific guidance and constructive feedback for the manuscript; Eric Hebert, Sukru Kaymakcalan, and Mark Aquino for their informatics support; and the antibody production group for generating the materials.

## Data availability statement

The datasets were not released due to confidential information. Example training script was provided in supplementary code.

