## Supplementary figures and images for "Protein language models enable prediction of polyreactivity of monospecific, bispecific, and heavy-chain-only antibodies"

### Supp Figure 1

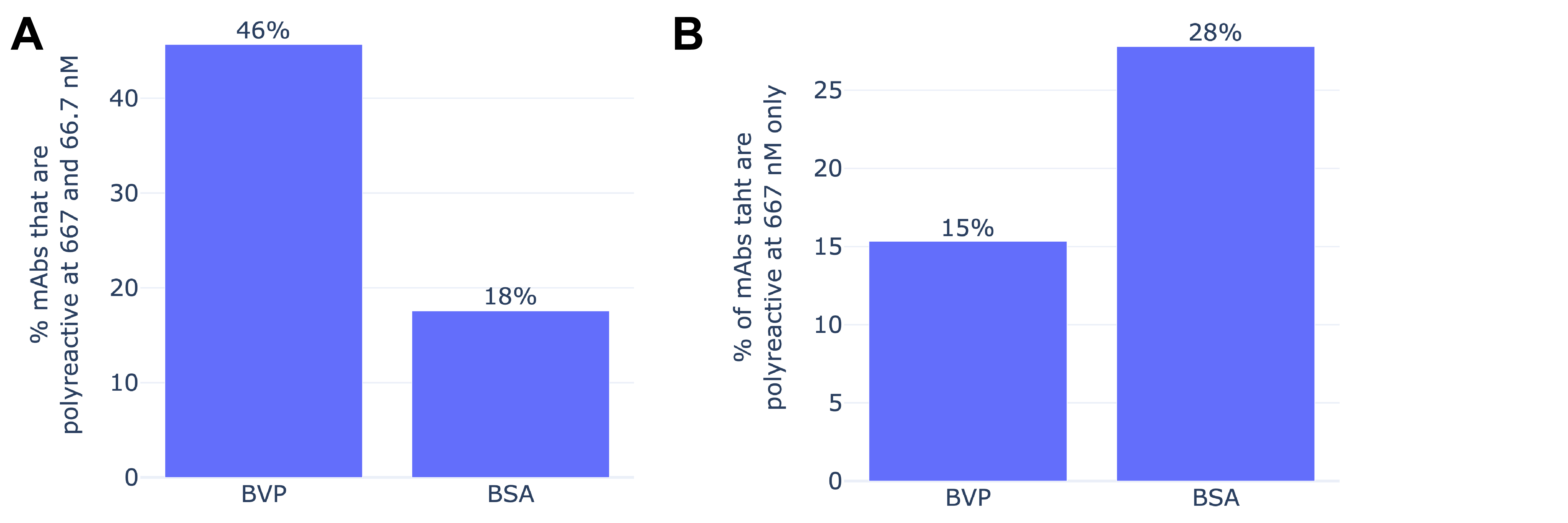

### Supp Figure 2

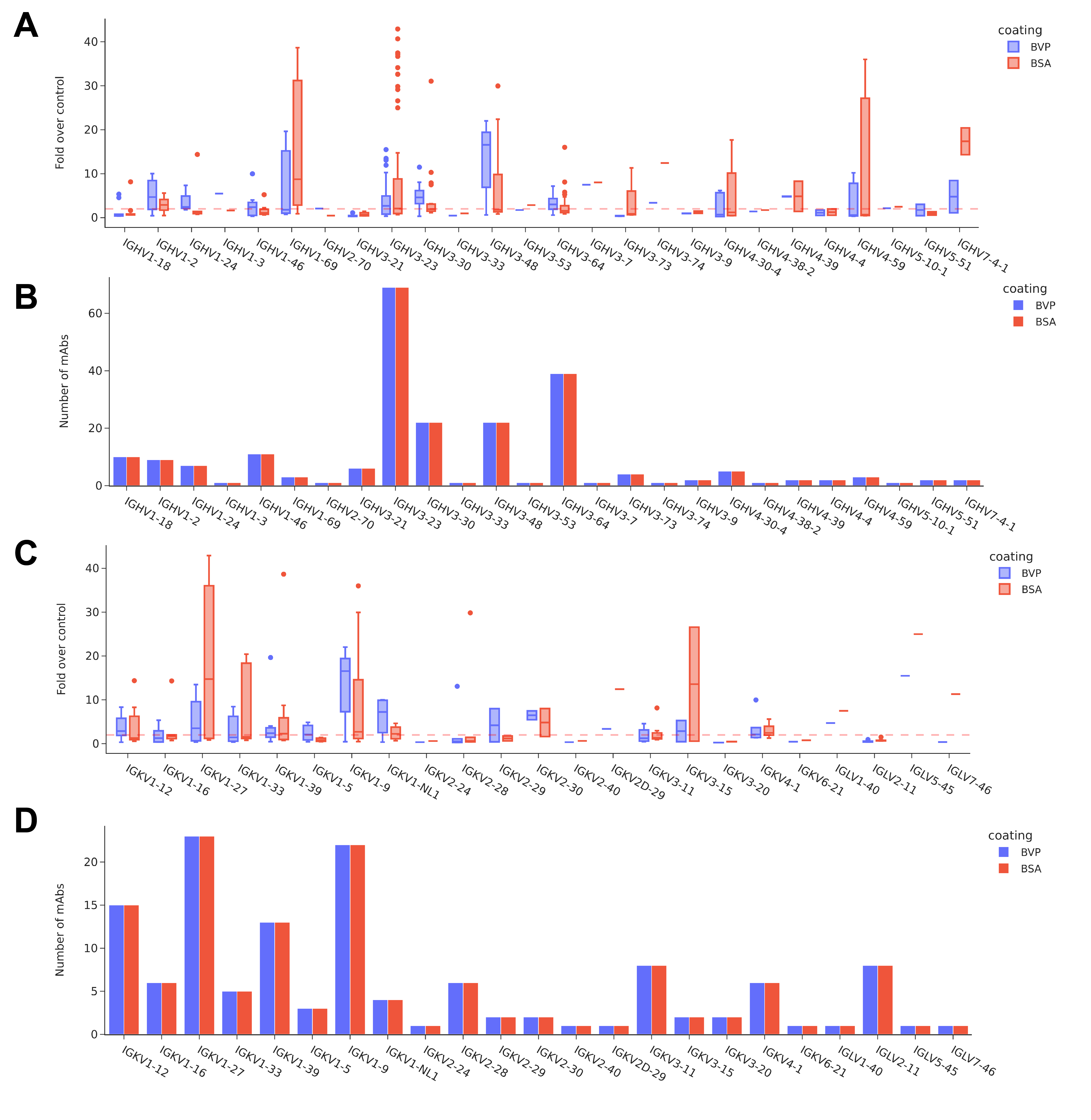

### Supp Figure 3

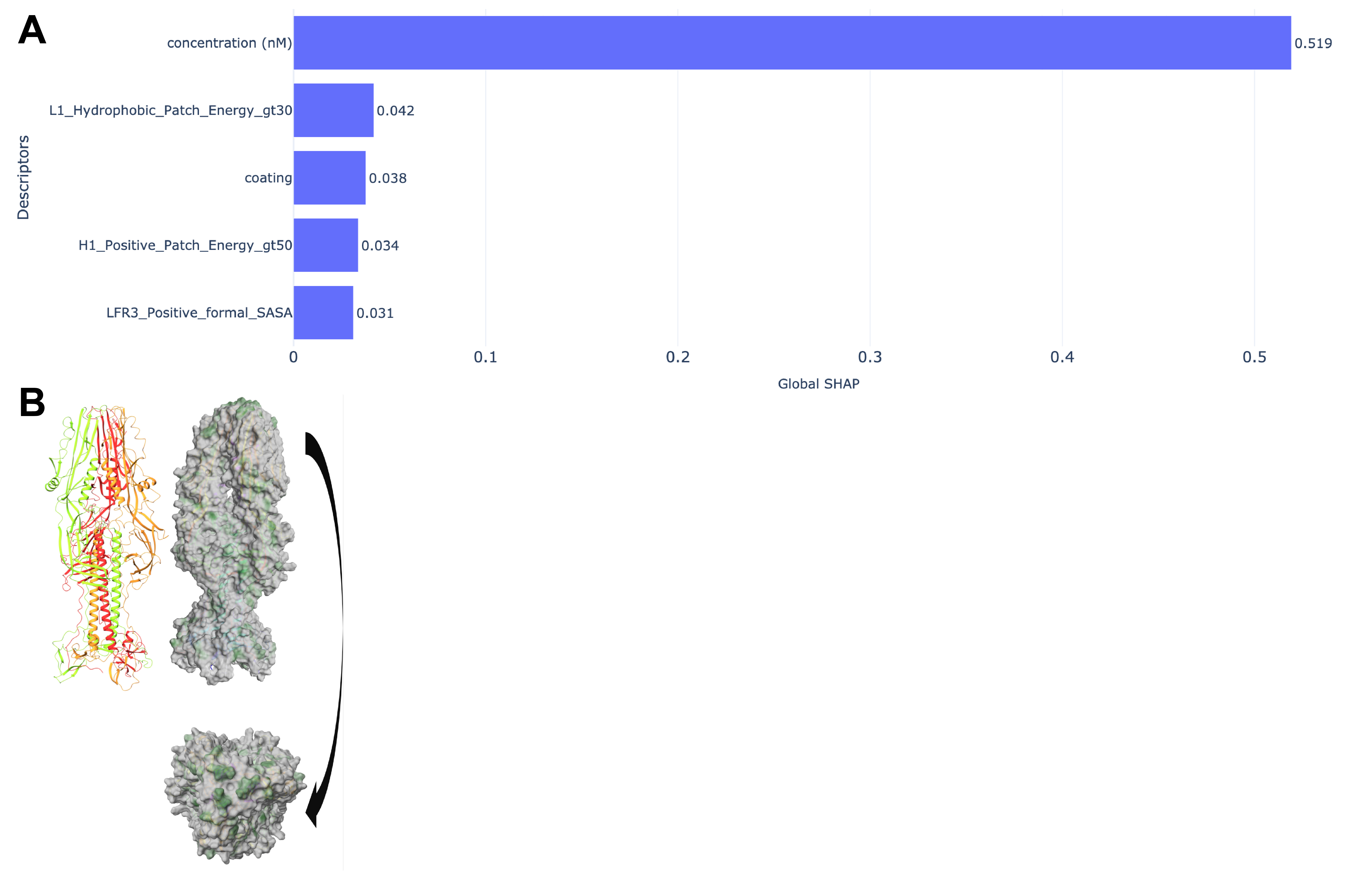
