## Supplementary material for "Protein language models enable prediction of polyreactivity of monospecific, bispecific, and heavy-chain-only antibodies": Supp Figure and Table Legends

Supp. Table 1. Performance of PLM models and ensemble on subsets of the September dataset

| **Model** | **Subset^1^** | **Size** | **ROC AUC^2^** | **Accuracy** | **Precision** | **Recall** | **F1** |
| --- | --- | --- | --- | --- | --- | --- | --- |
| PLM Model 2 | mAb with only HC | 79 | 0.841 | 0.797 | 0.741 | 0.690 | 0.714 |
|  | mAb with HC and LC | 521 | 0.876 | 0.787 | 0.773 | 0.776 | 0.775 |
|  | monospecific mAb | 480 | 0.873 | 0.802 | 0.765 | 0.784 | 0.774 |
|  | bispecific mAb | 120 | **0.862** | 0.733 | 0.787 | 0.716 | 0.750 |
|  | all (val+test) | 600 | 0.873 | 0.788 | 0.770 | 0.767 | 0.769 |
| PLM Model 4 | mAb with only HC | 79 | **0.968** | 0.886 | 0.778 | 0.966 | 0.862 |
|  | mAb with HC and LC | 521 | 0.864 | 0.775 | 0.728 | 0.837 | 0.779 |
|  | monospecific mAb | 480 | 0.889 | 0.808 | 0.730 | 0.885 | 0.800 |
|  | bispecific mAb | 120 | 0.812 | 0.717 | 0.746 | 0.746 | 0.746 |
|  | all (val+test) | 600 | 0.877 | 0.790 | 0.734 | 0.851 | 0.788 |
| PLM Model 5 | mAb with only HC | 79 | 0.916 | 0.785 | 0.800 | 0.552 | 0.653 |
|  | mAb with HC and LC | 521 | 0.865 | 0.781 | 0.789 | 0.732 | 0.759 |
|  | monospecific mAb | 480 | 0.879 | 0.802 | 0.809 | 0.712 | 0.757 |
|  | bispecific mAb | 120 | 0.808 | 0.700 | 0.738 | 0.716 | 0.727 |
|  | all (val+test) | 600 | 0.870 | 0.782 | 0.790 | 0.713 | 0.750 |
| Ensemble | mAb with only HC | 79 | 0.937 | 0.848 | 0.793 | 0.793 | 0.793 |
|  | mAb with HC and LC | 521 | **0.878** | 0.795 | 0.773 | 0.801 | 0.786 |
|  | monospecific mAb | 480 | **0.890** | 0.815 | 0.758 | 0.841 | 0.797 |
|  | bispecific mAb | 120 | 0.848 | 0.742 | 0.781 | 0.746 | 0.763 |
|  | all (val+test) | 600 | **0.883** | 0.798 | 0.760 | 0.818 | 0.788 |

^1^Combined analysis of September dataset's validation set and test set. "All (val+test)" refers to this aggregate, with other subsets examining mAbs by HC-only (VHH-Fc), HC+LC, monospecific (normal IgG and monospecific VHH-Fc), or bispecific categories.

^2^Bold numbers indicate the best ROC AUC values in each subset.

Supp. Figure 1. Effect of mAb concentration on BVP and BSA ELISA. A total of 313 unique mAbs from July dataset were tested at both 667 nM and 66.7 nM. **A**: Percent of mAbs that showed polyreactivity (fold of signal over control greater than 2) in BVP and BSA ELISA at both concentrations. **B:** Percentage of mAbs that exhibited polyreactivity at 667 nM but appeared clean (fold of signal over control equal to or less than 2) at 66.7 nM.

Supp. Figure 2. Polyreactivity differences among allele families. **A** and **B**: Analysis of 228 unique monospecific mAbs (regular mAbs and VHH-Fc) tested at 667 nM in BVP and BSA ELISA. X-axis shows allele families. Y-axis in **A** shows fold over control, and in **B** shows number of mAbs per family. **C** and **D**: Analysis of 134 unique monospecific mAbs (regular mAbs only) tested at 667 nM in both assays. Axis labels are similar to **A** and **B**, except light chain families were shown instead.

Supp. Figure 3. Mechanistic insights. **A**: The five most important features in the descriptor model, as determined by the SHAP score. L1: LC CDR1. H1: HC CDR1. LFR3: LC FR3. Detailed information on the descriptors can be found on Schrodinger website. **B**: AlphaFold predicted structure (ribbons) and hydrophobic surfaces (green patches) for GP64 trimer. Views are shown looking sideways through the trimer surface (top) and down along the trimer surface (bottom).
